# In vivo MRI measurement of microstructural constraints for direct delivery of therapeutics within the brain

**DOI:** 10.1101/2025.04.15.648763

**Authors:** Antonella Castellano, Valentina Pieri, Alice Segato, Marco Riva, Nicolò Pecco, Tian Yuan, Marco Vidotto, Davide Danilo Zani, Stefano Brizzola, Marco Trovatelli, Giuliano Ravasio, Riccardo Secoli, Stefano Galvan, Daniele Dini, Lorenzo Bello, Ferdinando Rodriguez y Baena, Elena De Momi, Andrea Falini

**Affiliations:** Vita-Salute San Raffaele University, Milan, Italy; Neuroradiology Unit and CERMAC, IRCCS Ospedale San Raffaele, Milan, Italy; Department of Electronics, Information and Bioengineering, Politecnico di Milano, Milan, Italy; Department of Biomedical Sciences, Humanitas University, Pieve Emanuele, Milan, Italy; IRCCS Humanitas Research Hospital, Rozzano, Milan, Italy; Department of Mechanical Engineering, Imperial College London, South Kensington Campus, Exhibition Road, London, UK; Department of Veterinary Medicine and Animal Sciences (DIVAS), Università degli Studi di Milano, Milan, Italy; S.S.D. Banca dei Tessuti e Terapia Tissutale, Grande Ospedale Metropolitano Niguarda, Milan, Italy; Department of Oncology and Hemato-Oncology, Università degli Studi di Milano, Milan, Italy; IRCCS Ospedale Galeazzi-Sant’Ambrogio, Milan, Italy

## Abstract

Brain tissue microstructure influences the efficient delivery of therapeutics within the brain. Diffusion Tensor Imaging (DTI) enables the depiction of tissue properties in vivo, and thus is potentially relevant for planning convection-enhanced delivery (CED) within the brain. We report on the quantitative assessment of the distribution of a Gadolinium solution infused by CED within the brain of a live ovine model. Infusate distributions were measured at multiple timepoints and compared to microstructural properties as depicted by DTI, thus demonstrating the impact of tissue features and catheter positioning on drug distribution in vivo. This study contributed to the clinical translation of the CED for flow-based therapy to ultimately provide new therapeutic approaches for several brain diseases, by providing essential tools and results used to develop a better prediction model and a delivery platform to reach the therapeutic target more precisely and non-invasively.

## Introduction

Convection-enhanced delivery (CED) of therapeutics to the brain is an approach for local delivery of personalized treatments directly into the parenchyma, overcoming the limited tissue-penetration of most drugs after systemic administration (1, 2). By generating a pressure-gradient at the tip of specialized infusion catheters, CED allows a targeted drug release that effectively bypasses the highly selective blood-brain barrier (BBB), irrespectively from the polarity or molecular weight of compounds (3, 4). Despite its wide-ranging applicability in different neurodegenerative diseases (5, 6), genetic disorders of metabolism (7), epilepsy, and brain tumors (8, 9), and its suitability for delivering different chemotherapies (10, 11), small molecules (12), engineered-toxins (13, 14), antiseizure agents, and gene therapies (15, 16), confirmatory evidence of CED efficacy in clinical trials is still limited (17).

Besides drug-specific pharmacological issues, several CED-specific physical limitations emerged, including backflow, inadequate target-coverage, unsuitable protocols, and devices, haphazard catheter positioning and inaccurate prediction of drug volume distribution (2, 18). The latter is such a pivotal issue for safe and effective pre-operative planning, that many computational models have been developed to simulate CED infusions and refine their predictive capacity (19–21). Drug distribution is known to be affected by infusion rate, catheter geometry, and flow of brain fluids – i.e., cerebrospinal fluid (CSF)/blood/interstitial fluid (22, 23). The influence of underlying tissue microstructure can play a crucial role in this context, as white matter spatial heterogeneity determines differential hydraulic permeability to flow, as recently demonstrated with post-mortem evaluations(24). State-of-the-art computational models included estimates of brain tissue anisotropy and diffusion non-Gaussianity to obtain more reliable predictions of infusate distribution volumes (25–28), which resulted in accurate matching between models and measurements performed in clinical experimental settings (26, 27). Nevertheless, it is still controversial whether tissue microstructure permeability, a fundamental parameter for modeling and deploying CED(28–32), play a role in vivo in the differential fluid flow in the brain parenchyma according to the different white matter (WM) spatial organization. It is still a matter of debate how to properly depict and to reliably quantify these phenomena non-invasively.

The non-invasive method commonly exploited to indirectly probe brain tissue properties in vivo is Diffusion Tensor Imaging (DTI), which derives a diffusion tensor ellipsoid from each voxel of diffusion-weighted Magnetic Resonance Imaging (MRI). The direction of the ellipsoid’s longest axis, along which water molecules preferentially move, is mathematically defined as the principal eigenvector (ε_1_) and corresponds to the major orientation of the axonal fibers in the WM (33). DTI has been previously exploited in simulation studies for guiding and planning CED procedures in human brain (34–36). Nonetheless, its full potential remains underexplored, particularly in both clinical studies on human subjects and most preclinical studies involving animal models such as swine (37), monkeys (38), sheep (39) and rats (40). Notable exceptions include studies that have used tractography reconstructions to target specific brain structure(41), or investigated the relationship between tissue properties and infusate distribution (42, 43). Despite these advancements, there is still a clear need to develop a fully integrated in vivo analysis incorporating a pre-operative CED planning tailored by DTI and tractography and a postoperative evaluation of brain microstructural constraints based on DTI. Such an approach will be able to provide missing patient-specific information, which can be used to influence drug distribution over time and prove previous experimental findings, where the WM spatial heterogenous organization has been shown to influence a differential flow (24). A precise characterization of the brain microstructure at the exact infusion site would be valuable for enhancing our understanding and potentially predicting the actual spatial distribution of a drug in an individual patient(44).

In this context, this study aimed at integrating DTI imaging studies into a new technology platform for minimally invasive and high precision drug delivery and testing it in the ovine model in vivo. It is hypothesized that the main direction of the WM fiber bundle, in related to direction of the flow, will influence final drug distribution. The main goal was to precisely characterize the drug distribution in the brain tissue after CED in vivo:

– by defining the microstructural properties of the ovine brain tissue using DTI.
– by tuning CED preoperative planning in sheep according to DTI-based tractography reconstructions, to infuse a gadolinium solution.
– by assessing the impact of brain microstructural features, as depicted by DTI, on drug distribution, facilitating it in any direction.
– by studying drug distribution after infusions both parallel and orthogonal to white matter fiber tracts to provide quantitative measures on how different orientations of the catheter delivering the infusion may influence CED.

## Materials and methods

### Study population and ethics

Seven adult female sheep (ovis aries, mean weight 72.2 ± 5.4kg) were included in this study, carried on in the context of the EU’s Horizon EDEN2020 project (https://www.eden2020.eu/). All animals were treated according to the European Communities Council directive (2010/63/EU), to the laws and regulations on animal welfare enclosed in D.L.G.S.26/2014 and in accordance with ARRIVE guidelines. Ethical approval for this study was obtained by the Italian Health Department with authorization n 635/2017.

### Anesthesia and MR Imaging Acquisition

All sheep were anesthetized via intravenous administration of Diazepam (0.25mg/Kg) + Ketamine (5mg/Kg), intubated and then maintained under general anesthesia with isoflurane (2%) and oxygen (2L/min). Animals were placed in prone position on a 1.5T clinical scanner (Achieva, Philips Healthcare) in a veterinary imaging facility [Fondazione La Cittadina Studi e Ricerche Veterinarie, Romanengo (CR), Italy] and MR imaging was performed to guide the neurosurgical planning for CED procedures. Small and medium flex coils were fixed over both ovine brain hemispheres. Presurgical T1-weighted volumetric scan was acquired from each animal by using a three-dimensional fast-field-echo sequence (3D-T1 FFE) (Figure 1A) with the following parameters: TR/TE 25ms/5ms; flip angle 40°; voxel size 0.667 × 0.667 × 1.4mm; SENSE factor R=2; 150 slices; acquisition time 8min 40s. Diffusion Tensor Imaging (DTI), then, was obtained by using a single-shot echo planar sequence with parallel imaging (SENSE factor R=2). Diffusion gradients were applied along 15 non-collinear directions, using a b-value of 1000s/mm^2^. The detailed imaging parameters for DTI were TR/TE 6,700 ms/84ms; acquisition isotropic voxel size 2 × 2 × 2mm; acquisition matrix 96 × 96; FOV 192 × 192mm; slice thickness 2mm; 45 contiguous slices without gap. Two signal averages (NSA =2) were obtained, for a total scan time of 5 min 34s. All the MRI sequences were oriented perpendicular to the longitudinal axis of the scanner without rotation in any plane, in order to minimize the requested steps for coregistration.

**Figure 1.**
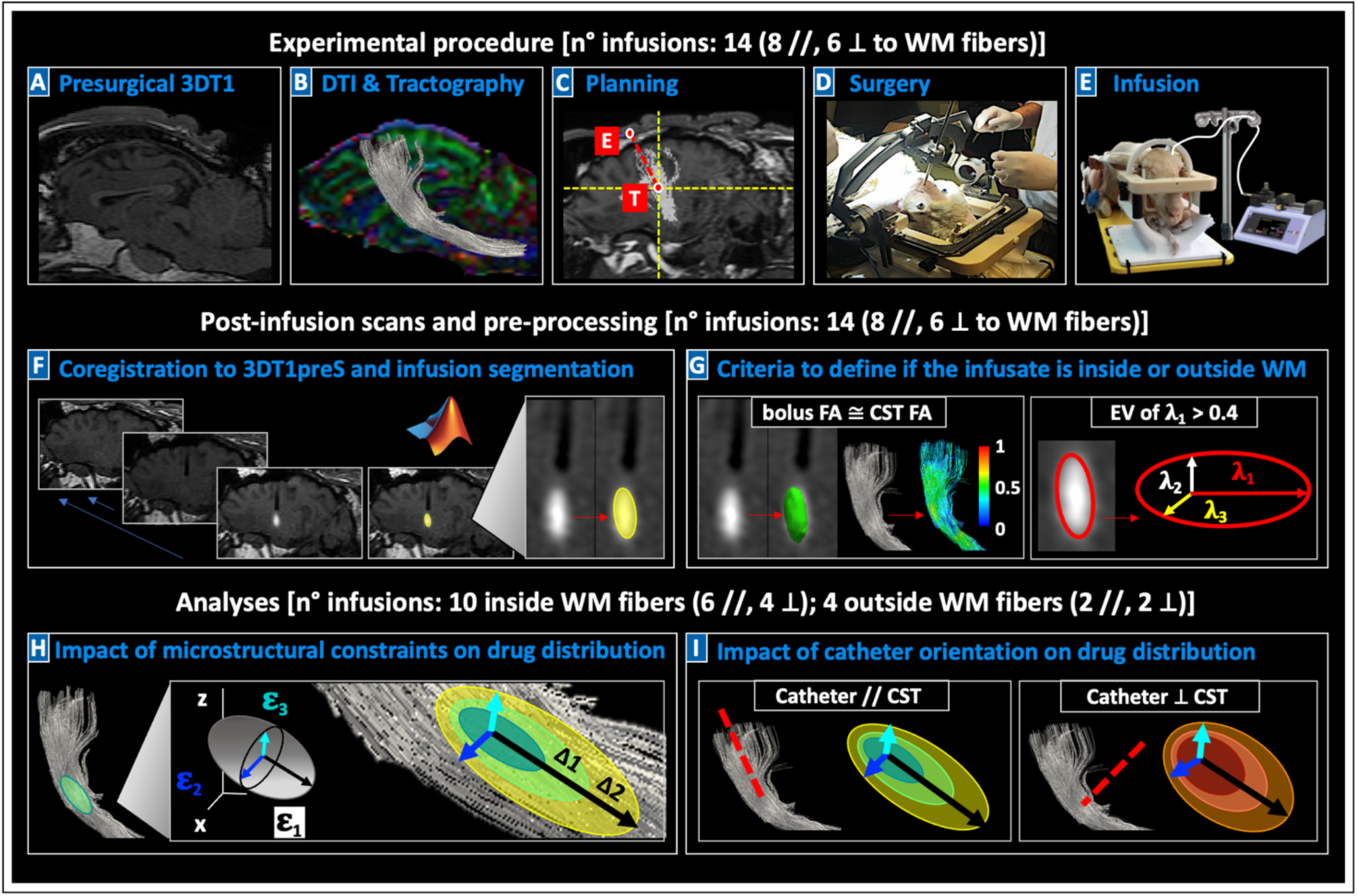
Workflow. Representation of the steps necessary for the experimental procedure [A-E], image pre-processing [F-G] and analyses [H-I]. **A)** For each sheep, first of all, a T1w imaging study was performed in order to exclude anatomical abnormalities of the brain. **B)** DTI was acquired (b=1000 s/mm^2^; 15 directions) and processed, allowing the reconstruction of CSTs. **C)** Pre-surgical planning consisted in the identification of a target (T) and entry (E) point to guide the catheter insertion, exploiting the T1 sequence on which the CST has been coregistered and superimposed. **D)** Catheters were stereotactically inserted by surgeons. **E)** 10 μL of a gadolinium-based solution were infused at 3 μL/min. **F)** Post-infusion T1w imaging study was performed immediately after stopping CED and at consecutive TP; the T1 for all the TP were coregistered by FSL, boluses were segmented with Matlab. **G)** FA of boluses and their EV were calculated, to establish if infusions reached the expected target within the WM, or ended up partially outside WM. **H)** Microstructural properties of brain tissue were correlated to distribution of infusions within brain tissue. **I)** Infusions performed with the catheter parallel to CST were compared to those performed with the catheter orthogonal to CST.

### Experimental procedure: pre-operative planning and infusions

After eddy-current correction by FMRIB Software Library (FSL, University of Oxford, https://fsl.fmrib.ox.ac.uk/fsl/), diffusion tensor and FA map were estimated. Ovine whole brain deterministic tractography was performed using the Diffusion ToolKit software (www.trackvis.org/dtk), that reconstructed fibers by a continuous tracking algorithm, keeping an FA threshold of 0.15 and an angle threshold of 27° as stopping parameters for the algorithm(45). Then, ovine corticospinal tracts (CSTs) were extracted from the whole brain tractography using TrackVis software (www.trackvis.org), by manually defining inclusion ROIs at the level of pons and internal capsule on the conventional color-coded FA maps(46), on the basis of a recently published ovine tractography atlas(45)(Figure 1B). Each CST was coregistered and superimposed to the 3DT1-FFE reference image of the same sheep and loaded on a bespoke version of the Neuroinspire^TM^ neurosurgical planning software (Renishaw plc.) (Figure 1C). The internal capsule was taken as target (T) point for the infusions, an entry (E) point was appropriately defined (Figure 1C), and a guide-catheter was inserted toward T via a bespoke MRI-compatible ovine headframe(47) and CRW stereotactic system (Figure 1D). The guide-catheter had an outer diameter of 2.5 mm and an internal lumen, to allow the insertion of an infusion catheter (fused silica of 238 μm diameter - Polymicro Nanocapillary TSP150238). Placement of the guide catheter was optimized according to the main direction of CST fibers in the target point, aiming at reaching CST fibers from a parallel orientation in eight cases, from an orthogonal orientation in six. To exclude hemorrhages or complications induced by surgery, animals underwent a 3DT1-FFE baseline MR acquisitions after the guide-catheter insertion. Then, sheep were removed from the scanner and the specialized infusion silica catheter for CED was inserted through the internal hollow of the guide catheter. A syringe pump (Pump 11 Elite & Pico Plus, Harvard Apparatus, Holliston, Massachusetts, USA) was connected to the infusion silica catheter through a dedicated extension line (1.5 m length, 1 mm diameter), in order to deliver a total of 10 μL of a gadolinium-based solution (Prohance®, 1:80 in saline) into each internal capsule. CED infusion rate was set at 3 μL/min, leading to a total infusion time of 3 minutes and 33 seconds for each infusate bolus (Figure 1E).

### Post-infusion scans and Pre-processing

Post-infusion 3DT1-FFE scans were acquired after stopping CED, at multiple timepoints (TPs). Consecutive images were obtained between 10 and 120 minutes after infusions, from a minimum of one scan to a maximum four scans, for each sheep. Specifications on the fourteen infusions are reported in Table 1. Post-infusion 3DT1-FFE scans of each animal were coregistered to the 3D-T1FFE baseline acquisition performed just after the guide-catheter insertion. Imaging values were normalized to an homogeneous intensity region in the temporal muscle, by means of FSL, to consistently estimate the differences in intra-sheep gadolinium intensities between consecutive TPs scans. Binary masks of the infused boluses for all TPs were extracted by an automatic region-growing algorithm (Matlab R2019a, Mathworks inc. Refer to Supplementary Material). The algorithm took the maximum gadolinium intensity of each bolus at the first TP scan as an input to calculate the maximum growth threshold, and then to limit the segmentation edges (Figure 1F). This procedure guaranteed homogeneous data handling and segmentation of gadolinium volumes across different TPs and animals. Segmentations were double-checked by a board-certified neuroradiologist with specific expertise in advanced clinical and preclinical imaging. Two criteria were applied to define whether the infusates ended up inside or partially outside WM, and to ensure reproducible analyses. First, average FA values were computed for all CSTs (*^CST^*FA*_mean_*) and for each bolus (*^bolus^*FA*_mean_*). Infusions were deemed inside WM fibers if their *^bolus^*FA*_mean_* fall in the range of *^CST^*FA*_mean_* ± 2SD, otherwise they were considered partially outside WM. Then, the Explained Variance (EV) of the principal component of each bolus (**λ**_1_) was calculated, and only infusions exceeding the cutoff of EV_**λ**1_ > 0.4 were regarded as inside WM fibers (Figure 1G). Further details of those criteria are described in Supplementary Material.

**Table 1:**
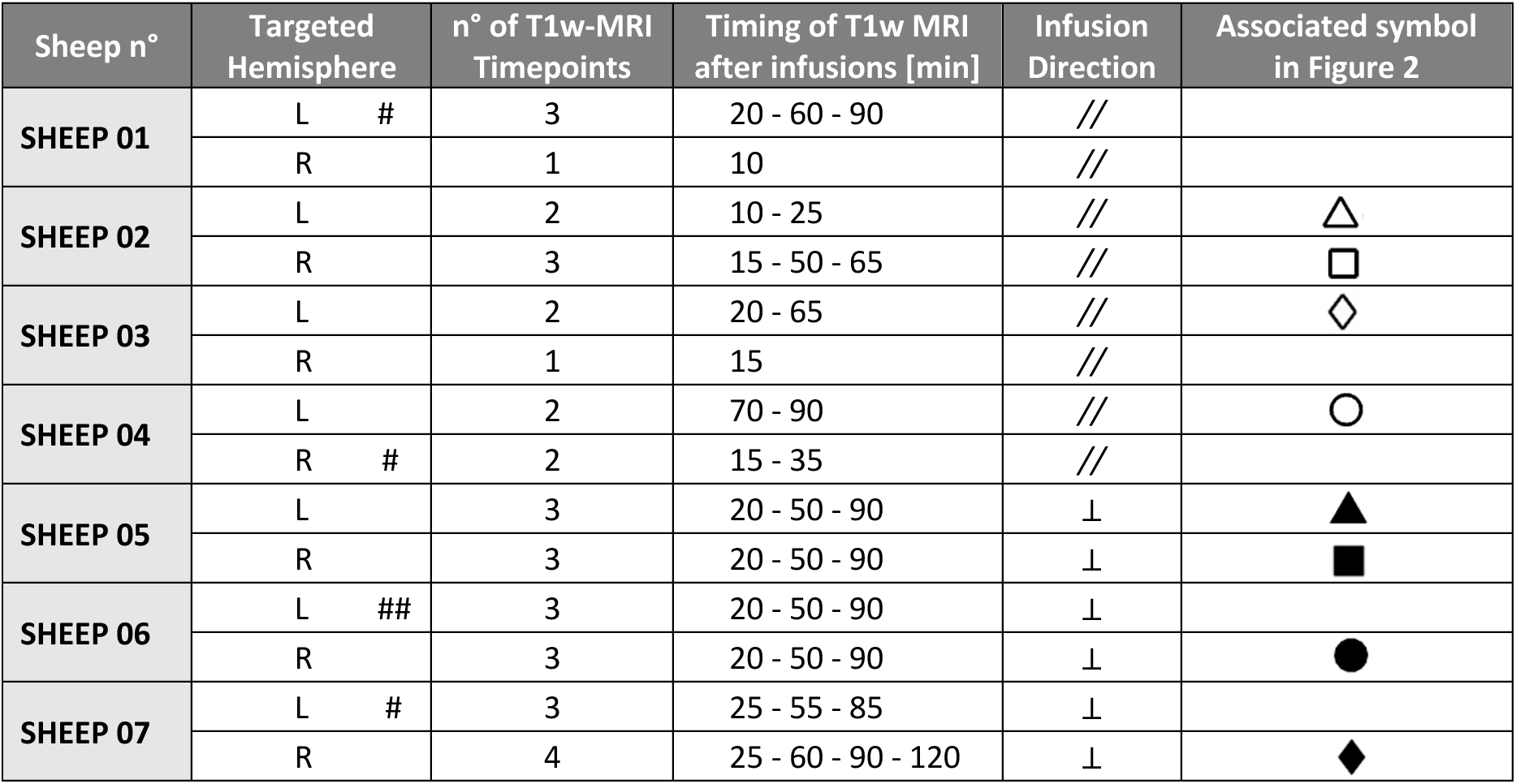
Specifications on the fourteen infusions in the ovine models. Details about the number of timepoints and the time-intervals between them is reported for each procedure. The direction of the catheter with respect to the WM of CST fibers was parallel (//) for eight infusions, orthogonal (⊥) for six infusions. Hashes indicate infusions outside WM fibers, as defined by the FA*_mean_* (#) and the EV (##) criteria. L= infusion in the left hemisphere, R= infusion in the right one.

| Sheep n° | Targeted Hemisphere | n° of T1w-MRI Timepoints | Timing of T1w MRI after infusions [min] | Infusion Direction | Associated symbol in Figure 2 |
| --- | --- | --- | --- | --- | --- |
| SHEEP 01 | L # | 3 | 20 - 60 - 90 | // |  |
|  | R | 1 | 10 | // |  |
| SHEEP 02 | L | 2 | 10 - 25 | // | △ |
| | R | 3 | 15 - 50 - 65 | // | □ |
| SHEEP 03 | L | 2 | 20 - 65 | // | $\diamond$ |
|  | R | 1 | 15 | // |  |
| SHEEP 04 | L | 2 | 70 - 90 | // | $\bigcirc$ |
|  | R # | 2 | 15 - 35 | // |  |
| SHEEP 05 | L | 3 | 20 - 50 - 90 | $\perp$ | ▲ |
| | R | 3 | 20 - 50 - 90 | $\perp$ | ■ |
| SHEEP 06 | L ## | 3 | 20 - 50 - 90 | $\perp$ | |
| | R | 3 | 20 - 50 - 90 | $\perp$ | $\bullet$ |
| SHEEP 07 | L # | 3 | 25 - 55 - 85 | $\perp$ | |
| | R | 4 | 25 - 60 - 90 - 120 | $\perp$ | $\blacklozenge$ |

### Post-infusion image analyses

As reported in Algorithm 1 (Refer to Supplementary Material) was exploited to compute the principal component analysis (PCA) relative to the maximal direction of each bolus across consecutive TPs (ε_max_bolus) and to the guide-catheter direction (Cath_direction), as well as to extract the DTI-derived eigenvectors (ε_1_, ε_2_, ε_3_) in each bolus location at the last TPs. Eigenvectors were weighted for the same-location FA values, to assign greater importance to voxels within WM. Those parameters were the basis for further estimates (Table 2A). Accordingly, bolus lengths (L) along ε_1_, ε_2_, ε_3_ were computed for each TP, and differences in L (ΔL) between the last TPs and the first one were calculated. ΔL_ε_1_, ΔL_ε_2_ and ΔL_ε_3_ of all the boluses inside WM fibers allowed us to study different infusate elongations along the three DTI-derived eigenvectors, evaluating the impact of microstructural constraints on drug distribution (Figure 1H, Table 2B).

**Table 2:** Parameters extracted from DTI images, for each bolus location. **A)** Basal parameters extracted from DTI images by means of FSL and 3DSlicer software, through python script. Cath_direction, ε_1_, ε_2_ and ε_3_ are constant, while ε_max_bolus changes for every timepoint. All of these parameters are 3D vectors. **B)** Combination of parameters allowed the calculation of the elongation and directionality of each bolus over time, along the DTI principal components. Therefore, they were distinguished depending on the direction (along ε_1_, ε_2_ and ε_3_).

| A | Basal Parameters | Description |
| --- | --- | --- |
| | $^{CST}FA_{mean}$ | Average FA of the CST fibers, calculated for all the fourteen infusions |
| | $^{bolus}FA_{mean}$ | Average FA of the infusate bolus, calculated for all the fourteen infusions |
| | EV | Explained Variance between the main eigenvalue of the bolus ($\lambda_1$) and its secondary eigenvalues ($\lambda_2, \lambda_3$) |
| | $\epsilon_{\text{max\_bolus}}$ | PCA relative to the maximal direction of the bolus |
|  | Cath_direction | PCA relative to the catheter direction |
| | $\epsilon_1$ | Primary eigenvector of the DTI voxels. In this study, it was extracted in correspondence to the location of the infusate bolus at the last timepoint acquired at MRI |
| | $\epsilon_2$ | Secondary eigenvector of the DTI voxels. In this study, it was extracted in correspondence to the location of the infusate bolus at the last timepoint acquired at MRI |
| | $\epsilon_3$ | Secondary and minimal eigenvector of the DTI voxels. In this study, it was extracted in correspondence to the location of the infusate bolus at the last timepoint acquired at MRI |
| B | Combination of Parameters | Description |
| | $L_{\epsilon_1}$ | Length of the bolus in the direction of $\epsilon_1$ [mm] |
| | $L_{\epsilon_2}$ | Length of the bolus in the direction of $\epsilon_2$ [mm] |
| | $L_{\epsilon_3}$ | Length of the bolus in the direction of $\epsilon_3$ [mm] |
| | $\Delta L_{\epsilon_1}$ | Difference in bolus length between consecutive timepoints and the first timepoint in the $\epsilon_1$ direction [mm]: bolus elongation along $\epsilon_1$ |
| | $\Delta L_{\epsilon_2}$ | Difference in bolus length between consecutive timepoints and the first timepoint in the $\epsilon_2$ direction [mm]: bolus elongation along $\epsilon_2$ |
| | $\Delta L_{\epsilon_3}$ | Difference in bolus length between consecutive timepoints and the first timepoint in the $\epsilon_3$ direction [mm]: bolus elongation along $\epsilon_3$ |
| | $\alpha$ | Angle between Cath direction and $\epsilon_1$ of CST fibers [°] |
| | $\Omega_{\epsilon_1}$ | Angle between $\epsilon_{\text{max\_bolus}}$ and $\epsilon_1$ [°] |
| | $\Omega_{\epsilon_2}$ | Angle between $\epsilon_{\text{max\_bolus}}$ and $\epsilon_2$ [°] |
| | $\Omega_{\epsilon_3}$ | Angle between $\epsilon_{\text{max\_bolus}}$ and $\epsilon_3$ [°] |

Then, angles (α) between Cath_direction and ε_1_, corresponding to the main direction of CST fibers in each bolus location, were calculated to check the infusion direction with respect to WM targets. Two groups were distinguished, so that infusions delivered parallel to CST could be compared to the orthogonal ones. EV of **λ**_1_ at first TP, that defined if boluses were inside WM, were used to quantitatively assess the shape of the infusates in parallel versus orthogonal infusions, immediately after stopping CED. Moreover, the length of boluses along the three main axes of DTI tensor (L_**ε**_1,_ L_**ε**_2_ and L_**ε**_3_) and the angle between ε_max_bolus and the three eigenvectors (**Ω**_**ε**_1_, **Ω**_**ε**_2_ and **Ω**_**ε**_3_) were used to evaluate the impact of catheter orientation on drug distribution over time, comparing the first with the last TPs (Figure 1I, Table 2B).

### Statistical analyses

Statistical analyses were achieved using GraphPad Prism v7 (GraphPad Software, San Diego, CA). Distributions of all variables were examined for normality using Shapiro-Wilk tests. Non-parametric Mann-Whitney U-test and Wilcoxon matched-pairs signed rank test were used for group comparisons. ANCOVA test was performed taking into account the time as a covariate and respecting its assumption(48). ΔL_ε_1_, ΔL_ε_2_ and ΔL_ε_3_ were analyzed through linear regression, and their slopes were compared through linear regression analysis tool (ANCOVA test). Boxplots described the distribution of data. Two-tailed p-value <0.05 was considered statistically significant.

## Results

The microstructural properties of the ovine brain tissue were consistently described by DTI in all animals, allowing reproducible tensor estimations and CST reconstructions with minimal inter-sheep variability. Pre-operative planning, surgical procedures and infusions were successfully accomplished in all the fourteen cases. Eight catheters were inserted parallel to CST fibers, six catheters orthogonal to them. We firstly calculated that three of the fourteen gadolinium-based boluses (two parallel and one orthogonal to CST) had a *^bolus^*FA*_mean_* lower than *^CST^*FA*_mean_* ± 2SD (#), and we secondly established that one bolus (orthogonal to CST) did not overcome the EV cutoff of 0.4, for its **λ**_1_ (##) (Table 1, Supplementary Material).

Accordingly, we acknowledged that four procedures did not reach the planned target and were delivered partially outside the WM; these trials were thus analyzed separately from the others. First, we considered all the infusions independently from the guide-catheter position, determining the influence of microstructural constraints of the brain tissue on drug distribution:

– inside WM fibers, as depicted by DTI, thus in an *anisotropic* environment (***3.1***, Figure 2);
– outside WM fibers, as depicted by DTI, thus in an *isotropic* environment (***3.2***, Figure 3);

**Figure 2.**
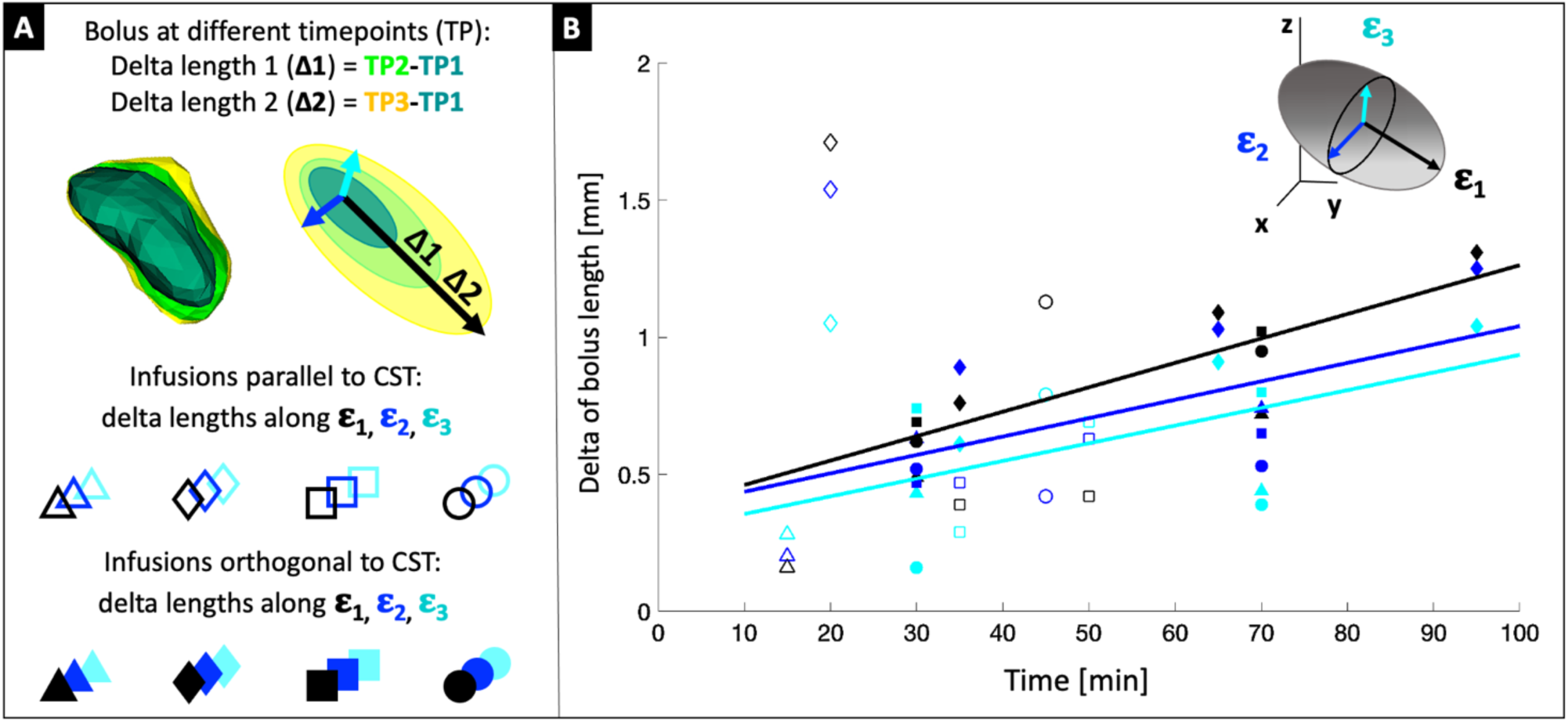
Impact of microstructural constraints on drug distribution inside WM fibers. The elongation of the shape of boluses inside WM fibers was studied over time, after stopping CED. Infusions delivered parallel and orthogonal to CST fibers have been studied together. **A)** Symbol legend, to interpret panel B. Delta length indicates the difference between consecutive timepoints (light green and yellow) and the first timepoint (dark green), and was measured along all the three main axes of DTI tensor. Each pair or triad of delta (ΔΩ_ε_1_, ΔΩ_ε_2_and ΔΩ_ε_3_) was plotted through the color-code relative to the three DTI directions (black= ε_1_, blue= ε_2_ and cyan= ε_3_), while different symbols were assigned to each sheep: empty for parallel infusions, color-filled for orthogonal infusions. **B)** Linear interpolations of each Δ along each direction: black line= ε_1_, blue line= ε_2_ and cyan line= ε_3_.

**Figure 3.**
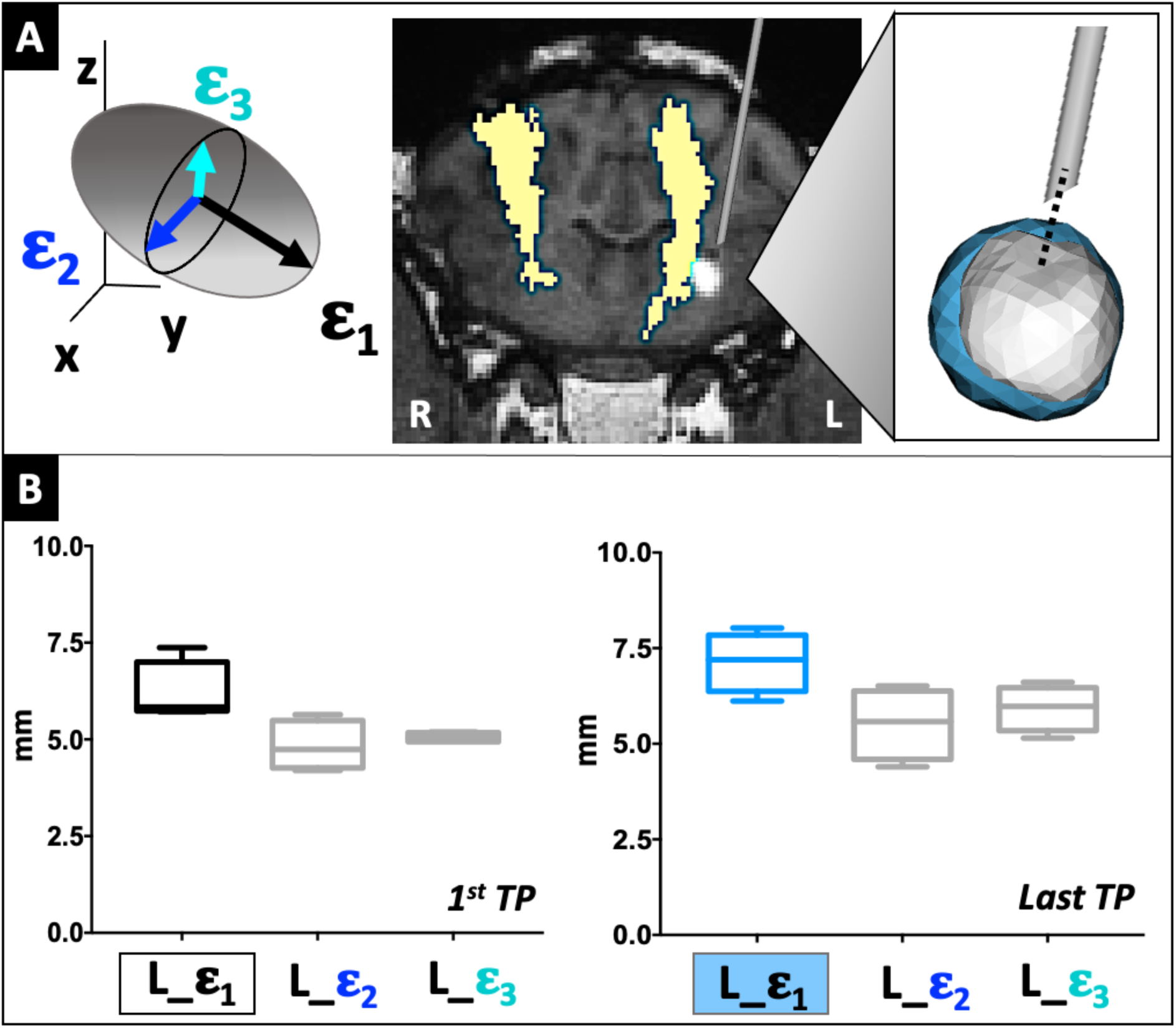
Impact of microstructural constraints on drug distribution outside WM fibers. The shape of the boluses infused outside WM fibers was studied over time, comparing their lengths along the three main axes of DTI tensor. **A)** Schematic representation of the tensor, highlighting its principal (black= ε_1_) and secondary (blue= ε_2_ and cyan= ε_3_), eigenvectors. Screenshot from FSL-eyes of a case-example of infusion outside WM (SHEEP 01, left side); in yellow, the binary mask of CST fibers coregistered and superimposed on the T1w image; the magnification shows the first (white) and last (light-blue) TPs of this infusion, that maintained a rounded shape over time. **B)** Boxplots showing the reciprocal relation between Ω_ε_1_, Ω_ε_2_and Ω_ε_3_ at the first (black-white and grey boxes) and last (light-blue and grey boxes) TP. Wilcoxon matched-pairs signed rank test did not highlight any significant difference between length of boluses along different DTI directions (P > 0.05), neither at the first, nor at the last TP.

We, then, focused exclusively on the infusions inside WM fibers, determining the influence of guide-catheter orientation with respect to WM fibers on drug distribution:

– at the first imaging TP after stopping CED, comparing EV and absolute L_ε_1_ of infusions parallel or orthogonal to CST fibers (***3.3***, Figure 4);
– over time, at consecutive imaging TPs, comparing the reciprocal evolution of L_ε_1_, L_ε_2_ and L_ε_3_ of infusions parallel or orthogonal to CST fibers (***3.4***, Figure 5);
– over time, at consecutive imaging TPs, comparing the reciprocal evolution of **Ω**_ε_1_, **Ω**_ε_2_ and **Ω**_ε_3_ of infusions parallel or orthogonal to CST fibers (***3.5***, Figure 6).

**Figure 4.**
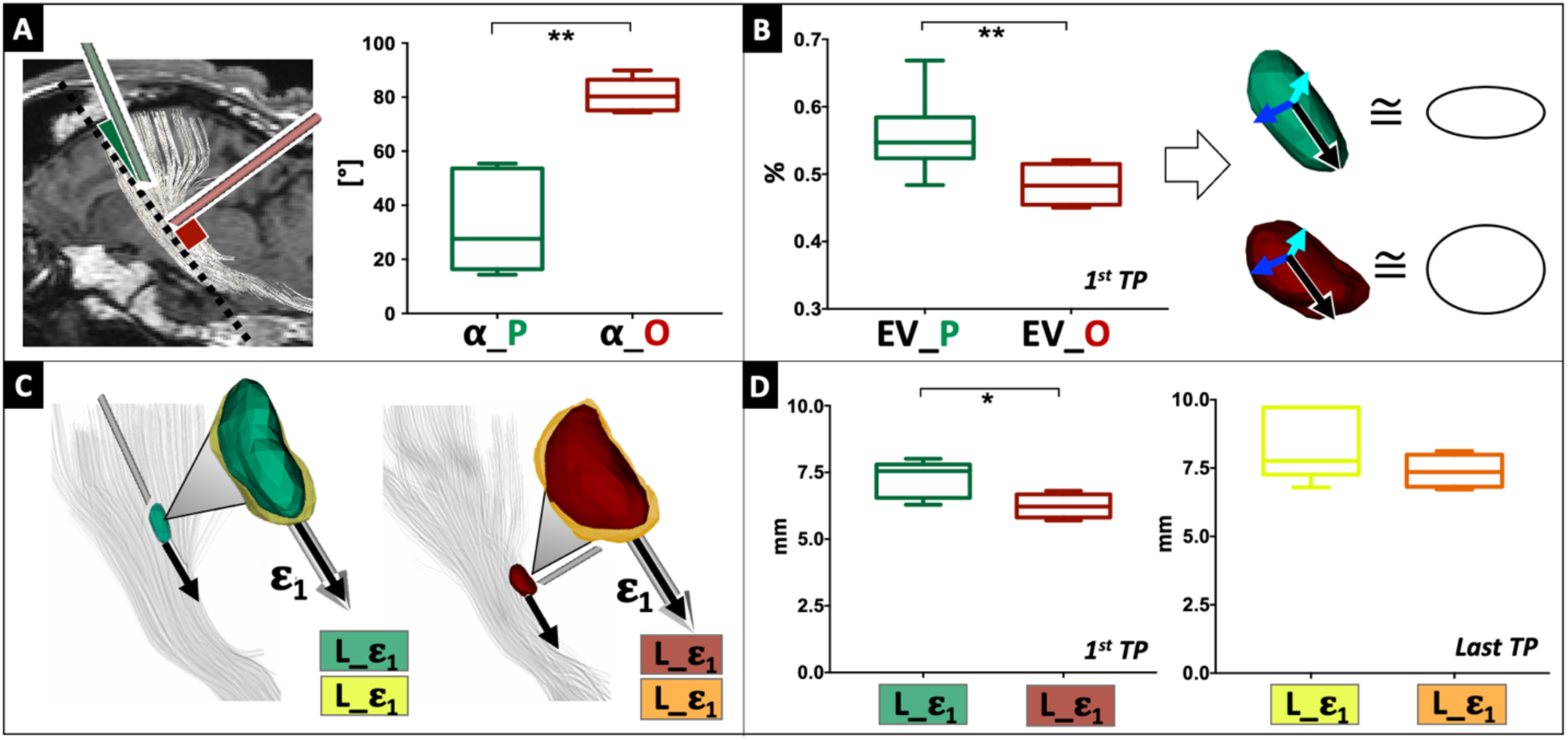
Impact of catheter orientation on drug distribution. Initial distribution of infusions inside WM fibers was studied, according to the parallel or orthogonal orientation of the catheter with respect to the CST (*P < 0.05, **P < 0.01). **A)** Angle **α** between the direction of CST fibers (dotted line) and catheters, depicted in green if the pre-surgical planning aimed at infusing parallel to WM, in red if the pre-surgical planning aimed at infusing orthogonal to it. Mann-Withney test confirmed clear difference between the six parallel and the four orthogonal infusions analyzed. **B)** Explained Variance (EV) of the boluses at the first time points, influenced by CED. Both parallel (green) and orthogonal (red) infusions had an oval shape, but ANCOVA test highlighted that the former were significantly more elongated than the latter. **C)** Symbol legend, to interpret panel D. **D)** Maximal lengths, along the main eigenvector, of boluses infused parallel to CST were significantly higher than their orthogonal counterparts at the 1^st^ MRI TP (green vs red, respectively), but this difference attenuated over time at the last MRI TP (yellow vs orange, respectively).

**Figure 5.**
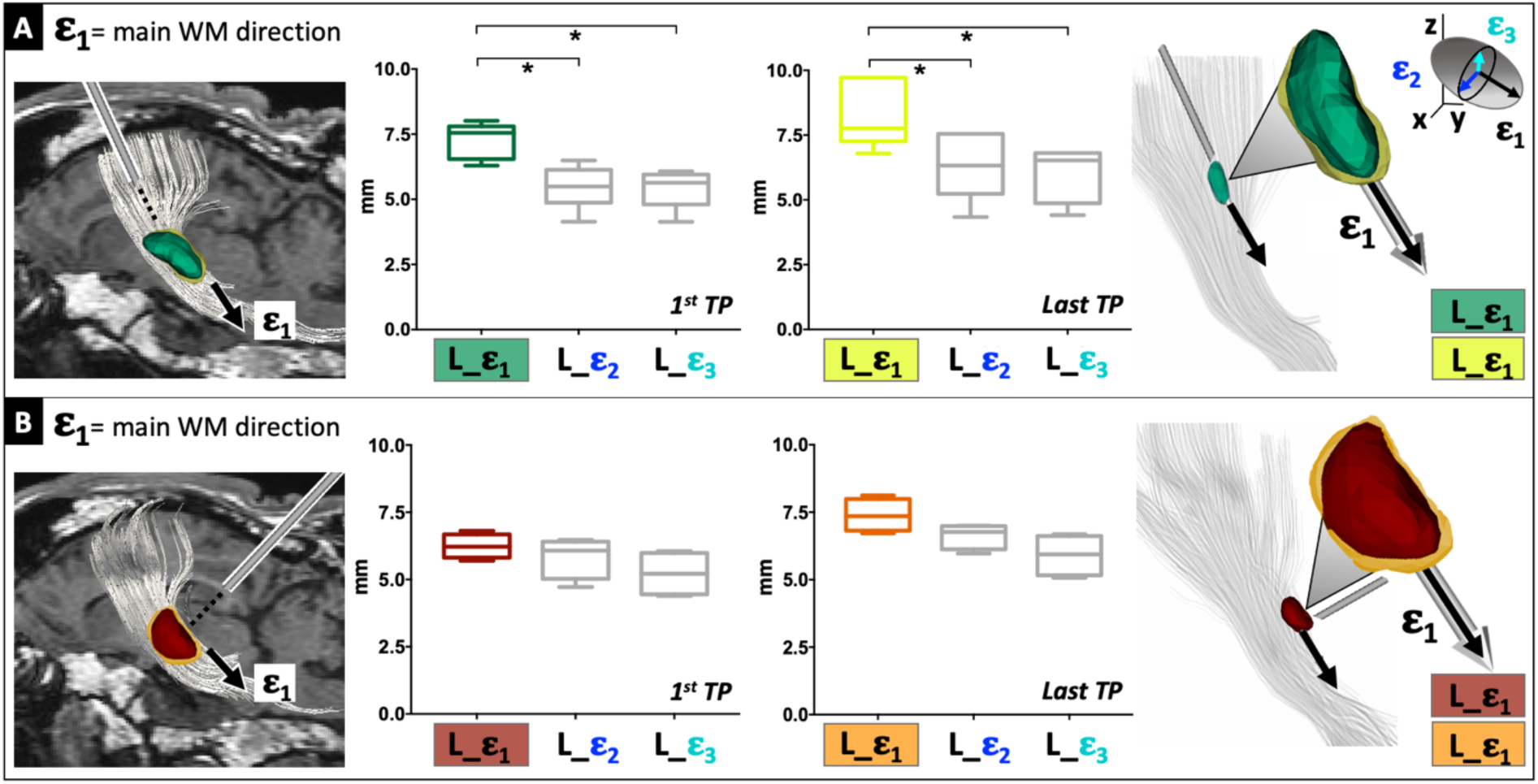
Impact of catheter orientation on drug distribution: lengths of the infusates inside WM fibers. Diffusion of infusions inside WM fibers over time was studied, according to the parallel or orthogonal orientation of the catheter with respect to the CST (*P < 0.05). Paired comparisons of bolus lengths (L) along the three main axes of DTI tensor (represented at the top right corner of the figure) were performed. **A)** Boluses infused parallel to CST fibers were evaluated at the 1^st^ and last TP of MRI after stopping CED. Significant differences between *L*_**ε**_1_, *L*_**ε**_2,_ and *L*_**ε**_3_ emerged both at the 1^st^ and last imaging TP. **B)** Boluses infused orthogonal to CST fibers were evaluated at the 1^st^ and last TP of MRI after stopping CED. Differences between *L*_**ε**_1_, *L*_**ε**_2_ and *L*_**ε**_3_ were not significant both at the 1^st^ and last imaging TP, but a trend towards the most prominent increase of Ω_**ε**_1_ over time was evident.

**Figure 6.**
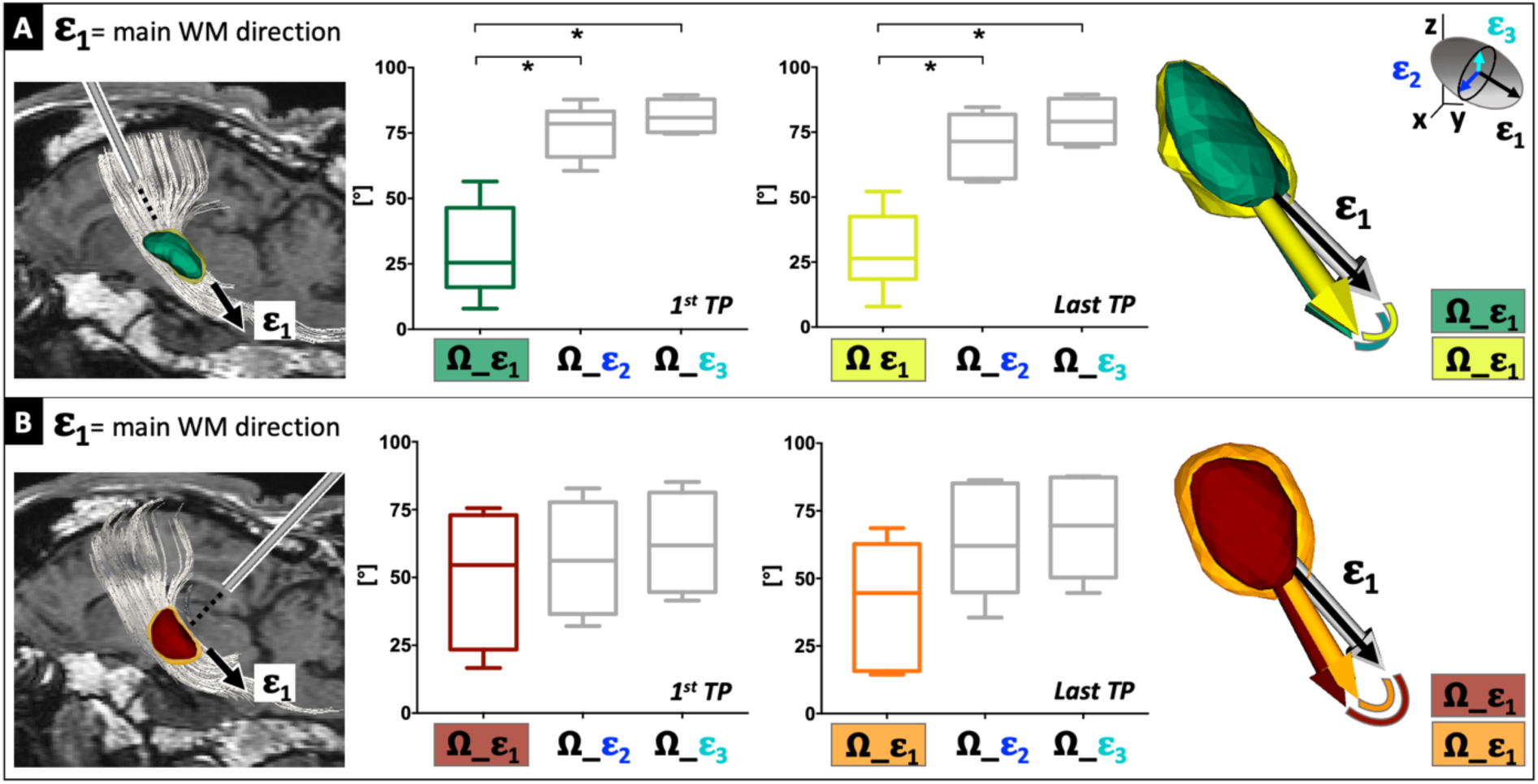
Impact of catheter orientation on drug distribution: angles between the infusates and WM fibers. Diffusion of infusions inside WM fibers over time was studied, according to the parallel or orthogonal orientation of the catheter with respect to the CST (*P < 0.05). Paired comparisons of angles (**Ω**) between the three main axes of DTI tensor (represented at the top right corner of the figure) and the PCA of the bolus was performed. **A)** Boluses infused parallel to CST fibers were evaluated at the 1^st^ and last TP of MRI after stopping CED. Significant differences between **Ω**_**ε**_1_, **Ω**_**ε**_2_ and **Ω**_**ε**_3_ emerged both at the 1^st^ and last imaging TP. **B)** Boluses infused orthogonal to CST fibers were evaluated at the 1^st^ and last TP of MRI after stopping CED. Differences between **Ω**_**ε**_1_, **Ω**_**ε**_2_ and **Ω**_**ε**_3_ were not significant both at the 1^st^ and last imaging TP, but a trend towards the most prominent decrease of **Ω**_**ε**_1_ over time was evident.

The following paragraphs illustrate results in detail.

### Infusions inside WM fibers: impact of brain microstructural constraints on drug distribution

The computation of ΔL, defined as the bolus elongation between consecutive TPs and the first TP along the DTI main directions (ε_1_, ε_2_ and ε_3_), allowed to investigate the influence of microstructural tissue features on drug distribution inside WM fibers, irrespectively from the direction of the guide catheter. Therefore, boluses derived from parallel and orthogonal infusions have been considered together, focusing on the shape-changes of each bolus among sequential measurements. Eight infusions were analyzed, since the ones checked at a single TP only were excluded from this study, as well as the ones partially outside WM fibers (Table 1, Supplementary Material). Resulting ΔL along ε_1_, ε_2_ and ε_3_ of every case, fitted by linear interpolations for each DTI direction, showed that the shape elongation of all boluses was maximal along the principal WM fiber direction. In fact, the slope of the line interpolating ΔΩ_ε_1_ was 0.5346 mm/h, greater than those interpolating ΔΩ_ε_2_ (0.4027 mm/h) and ΔΩ_ε_3_ (0.3871 mm/h) for all infusions, demonstrating the highest hydraulic conductivity along the main fiber axial direction ε_1_. Moreover, R^2^ value was 0.25 for ΔΩ_ε_1_, while it was 0.19 and 0.26 for ΔΩ_ε_2_and ΔΩ_ε_3_, respectively (Figure 2, Table 3).

**Table 3:**
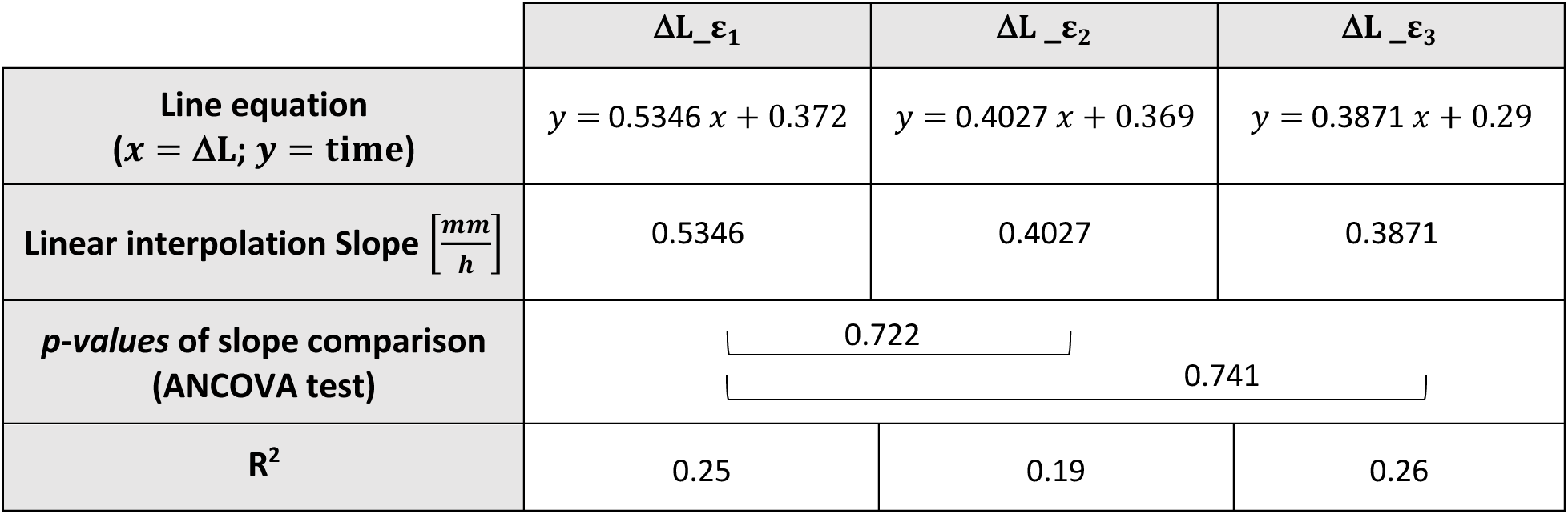
Quantitative analyses to define the impact of microstructural constraints on diffusion inside WM fibers. Linear interpolation slopes of ΔL _ε_1_, ΔL _ε_2_ and ΔL _ε_3_ indicated as [mm/h], graphically represented in Figure 2. ΔL _ε_1_ is greater than ΔL _ε_2_ and ΔL _ε_3_, but the ANCOVA test comparing the slopes was not statistically significant. R^2^ represents the coefficient of determination of the linear regression.

### Infusions outside WM fibers: impact of brain microstructural constraints on drug distribution

To characterize the time-evolution of the shape of four boluses resulting partially outside WM fibers, we analyzed their features at the first and last TP of MRI acquisitions. At the first imaging TP, length of those boluses along the main DTI eigenvector (L_ε_1_: median= 5.835 mm; range= [5.72-7.37 mm]) did not statistically differ from their length along the secondary DTI eigenvectors (L_ε_2_: median= 4.74 mm; range= [4.2-5.64 mm] and L_ε_3_: median= 5.035; range= [4.94-5.19 mm]), with a P > 0.05. The same result was obtained at the last imaging TP, after diffusion of the infusate in the brain tissue outside WM, where the difference between L_ε_1_ (median= 7.2 mm; range= [6.12-8.03 mm]), L_ε_2_ (median= 5.58 mm; range= [6.38-6.51 mm]) and L_ε_3_(median= 5.98 mm; range= [5.15-6.61 mm]) remained not statistically significant (P > 0.05) (Figure 3, Supplementary Table 2).

### Infusions parallel and orthogonal to WM fibers: impact of catheter orientation on EV and absolute L_ε_1_

The relevance of the catheter orientation to the CST was investigated through ten infusions inside WM fibers. The angle between catheters and fibers was computed, confirming that each actual catheter position matched the corresponding presurgical planning: six were parallel (median= 27.61°; range= [14.33-55.45°]), four orthogonal (median= 80.24°; range= [74.34-89.89°]) to CST fibers (Figure 4A). Being those angles statistically different between the two groups (P=0.0043), further comparative studies were performed. The initial analyses focused exclusively on the first TP of MRI acquisitions to highlight the differences of the boluses immediately after stopping CED.

EV at the first TP revealed that boluses delivered both parallel and orthogonal to CST had an oval shape (EV of **λ**_1_ > 0.4), but the former displayed higher EV (mean ± SD = 0.556 ± 0.60) than the latter (mean ± SD = 0.484 ± 0.31). Thus, infusions parallel to WM fibers resulted significantly more elongated along **λ**_1_ than orthogonal ones (P=0.008) (Figure 4B, Supplementary Table 3). In line with EV, at the first TP, L_ε_1_ of boluses infused parallel to CST was significantly longer than the L_ε_1_ of orthogonal infusions (P=0.0381) (Figure 4D), given a fixed quantity of gadolinium-based solution. Since the above-described difference in L_ε_1_did not remain significant at the last TP of MRI analyses (P > 0.05) (Figure 4D), we decided to further investigate the evolution of the shape of boluses over time, according to the different infusion orientations.

### Infusions parallel and orthogonal to WM fibers: impact of catheter orientation on L_ε_1_, L_ε_2_ and L_ε_3_

For infusions parallel to the CST fibers, L_**ε**_1_ (median= 7.55 mm; range= [6.29-8.01 mm]) was significantly greater than L_**ε**_2_ (median= 5.49 mm; range= [4.14-6.49 mm]) and L_**ε**_3_ (median= 5.63 mm; range= [4.13-6.07 mm]) already at the initial TP acquisitions (P=0.0312). This difference was maintained over time, remaining significant at the last TP, with L_**ε**_1_ (median= 7.76 mm, range= [6.79-9.72 mm]) longer than L_**ε**_2_ (median= 6.325 mm; range= [4.34-7.55 mm]) and L_**ε**_3_ (median= 6.525 mm; range= [4.41-6.81 mm]) (Figure 5A, Supplementary Table 3).

Conversely, infusions orthogonal to CST fibers showed a non-significant trend of a greater L_**ε**_1_ (median= 6.22 mm; range= [5.7-6.81 mm]) with respect to L_**ε**_2_ (median= 6.075 mm; range= [4.72-6.47 mm]) and L_**ε**_3_ (median= 5.215 mm; range= [4.39-6.05 mm]) at the first TP acquisitions (P > 0.05). Notably, this trend increased over time, with the last TP showing L_**ε**_1_ (median= 7.355 mm; range= [6.72-8.12 mm]) more detached from L_**ε**_2_ (median= 6.77 mm; range= [5.97-7 mm]) and L_**ε**_3_ (median= 5.935 mm; range= [5.07-6.68 mm]) (Figure 5B, Supplementary Table 3).

### Infusions parallel and orthogonal to WM fibers: impact of catheter orientation on **Ω**_ε_1_, **Ω**_ε_2_ and **Ω**_ε_3_

As far as **Ω** is concerned, the alignment of boluses to the WM fibers appeared consistent with the above-described results of L_ε_1_, L_ε_2_ and L_ε_3_. Specifically, **Ω**_**ε**_1_ of parallel infusions resulted significantly smaller than **Ω**_**ε**_2_ and **Ω**_**ε**_3_, both at the first imaging TP (**Ω**_**ε**_1_ median= 25.43°; range= [7.88-56.48°]; **Ω**_**ε**_2_ median= 78.56°; range= [60.56-87.77°]; **Ω**_**ε**_3_ median= 80.90°; range= [74.69-89.51°]), and at the last TP (**Ω**_**ε**_1_ median= 26.42°; range= [7.88-52.14°]; **Ω**_**ε**_2_ median= 71.44°; range= [55.9-84.63°]; **Ω**_**ε**_3_ median= 79.09°; range= [69.35-89.52°]) (P=0.0312) (Figure 6A, Supplementary Table 3). Finally, for infusions orthogonal to the CST fibers, a non-significant trend of reduced **Ω**_**ε**_1_ compared to **Ω**_**ε**_2_ and **Ω**_**ε**_3_ was observed at the first TP (**Ω**_**ε**_1_ median= 54.58°; range= [16.61-75.52°]; **Ω**_**ε**_2_ median= 56.17°; range= [32.36-82.82°]; **Ω**_**ε**_3_ median= 61.8°; range= [41.5-85.2°]) (P > 0.05). As it was observed in the length analysis, this trend strengthened at the last TP, where differences between **Ω**_**ε**_1_, **Ω**_**ε**_2_ and **Ω**_**ε**_3_ became more evident (**Ω**_**ε**_1_ median= 44.59°; range= [14.41-68.6°]; **Ω**_**ε**_2_ median= 62.01°; range= [35.5-86.34°]; **Ω**_**ε**_3_ median= 69.54°; range= [44.64-87.74°]) (Figure 6B, Supplementary Table 3).

## Discussion

Several factors contribute to the efficacy of the CED, such as catheter design and placement, tumor location and size, infusion rate and frequency, drug type and concentration, and brain anatomy. Among those, this study focused on the tissue microstructural constrains and the catheter pose that influence the distribution of a solution through the CED in vivo in a large animal model. This study also provided experimental evidence of tractography integration in CED pre-surgical planning. Results demonstrate that the subcortical white matter architecture, as depicted and quantitatively assessed by MR-DTI, is a pivotal factor shaping the solute distribution with the orientation of the infusion catheter along or perpendicular to the main axis of the WM fiber bundle. Fourteen CED procedures have been successfully accomplished in living sheep, and quantitative analyses of the infusates show that depicting tissue microstructure by DTI may be essential to plan optimal catheter insertion in individual patients, and to accurately predict the final drug distribution in CED.

### Tractography integration in presurgical planning for CED in ovine models

The individual presurgical planning was integrated with the sheep-specific DTI-based tractography, reconstructed for all animals before CED, to direct the infusions to the WM fibers with a higher degree of isotropy specifically. Tractography has been previously deployed in a similar pre-clinical scenario on macaques to simulate the intra-cerebral movement of molecules exclusively *a posteriori*, i.e. after placing the seed near the infusion target (41). This study deployed tractography to tune CED in vivo *a priori* for the first time, to the best of our knowledge. Our approach aimed at reproducing the actual clinical setting, where tractography is increasingly used as an integral part of both the pre-operative and intra-operative work-up for minimally invasive neurosurgical procedures. As an example in brain tumors, by visualizing the WM structures proximal to the peritumoral areas and possibly infiltrated by the mass, the surgeon has a valid imaging adjunct to maximize the removal of the diseased tissue while minimizing collateral damages to surrounding eloquent structures (49–51). A less explored scenario of application of tractography is the CED. CED may benefit from the integration of tractography in pre-operative planning since WM fibers may influence the delivery of therapeutics depending on the anisotropy of the targeted brain region. Previous data, in fact, showed that the brain tissue architecture shapes the fluid flow through the brain (52). This has major implications for drug delivery scenarios and additionally for physiological interstitial flow. As also further discussed below, the current results proved that a thorough pre-operative selection of the infusion target could accurately predict drug distribution volume over time and that MR-DTI.

### Influence of microstructural constraints of brain tissue on drug distribution

A further relevant issue of our study regarded the analysis of the microstructural properties of the ovine brain tissue in vivo with DTI, both within and outside the most compact bundles of WM.

On the one hand, we study drug distribution based on single-case DTI data, acquired in living ovine models, differently from a previous study that compared results of the in vivo CED in rats to a voxelized computational model built with DTI images acquired on fixed rat brains (42). On the other hand, we focused on how the drug distribution may vary depending on the final infusion target, within or outside the most compact portion of CST fibers, while Rosenbluth et al. performed serial DTI during CED in the thalamus, thus only in the gray matter (43).

For the infusions performed inside the WM fibers, thus in the anisotropic environment, quantitative analyses of ΔL along ε1, ε2, ε3 over time demonstrated that the microstructural constraints impact drug distribution, facilitating it along the main fiber direction (ε1). ΔL along ε1 was greater than ΔL along ε2 and ε3 in terms of absolute values, and it also keeps increasing more prominently over time. It is also important to highlight that the slope of the line interpolating ΔL_ε2 is steeper than that interpolating ΔL_ε3, further remarking how tensorial estimates may accurately depict brain tissue properties, thus gaining the potential to pilot microsurgical procedures.

For the infusions performed outside WM fibers, thus in an isotropic environment, a predominant direction of bolus elongation could not be determined, neither at the first TP after stopping CED nor over time. In fact, L_ε1 was not significantly greater than L_ε2 or L_ε3 at the first and last TPs, as it can also be appreciated by observing the rounded shape of the infusates. It should also be noted that the lengths along ε2 were always shorter than the measures along ε3; the infusate diffusion did not follow a preferential direction if the tissue is not markedly anisotropic, as outside the most compact bundle of the WM fibers.

These results demonstrated the infusate to spread along WM tracts preferentially rather than outside the WM bundles. While this phenomenon was as expected in parallel infusion cases, it was somewhat surprising in the orthogonal infusion cases, as it proves that WM microstructure dominates the response of the tissue and the drug distribution. This motivated a parallel study (53) in which we employed advanced computational fluid dynamics simulations to investigate this mechanism and found that resolving the mechanical interactions between the infused fluid and axons is key to investigate the diversion of the infusate along the axonal tracts in the conditions under investigation here.

This finding may be crucial if the impact of the fluid flow in brain tumors treatment is considered, including therapeutic strategies and drug transport. The dissemination of tumor cells along fibers to distant regions of the brain may be the main reason for the limited impact of current therapies on patients’ outcomes (52). In this regard, the preferential flow of interstitial fluid and drugs along the same paths may be exploited to finally reach those spreading malignant cells more efficiently (54). In addition to avoiding tumor dissemination, an improved understanding of drug distribution mechanisms through different brain areas is crucial to design more effective therapeutic approaches tailored on an individual basis according to the specific brain tissue features. This is particularly relevant in the context of CED procedures, effectively bypassing the BBB and allowing the local delivery of molecules larger than the standard chemotherapy drugs, such as immunotherapies and nanoparticles (54). Deploying clinical imaging, as MR-DTI, to identify and quantify the microstructural tissue constraints for volume distribution, the approach herein described probed that MR-DTI can be a reliable and reproducible method to depict these properties in a patient- and disease-specific manner. This could be essential to improve the understanding of patients’ pathological changes, especially in such a heterogeneous scenario as that of patients with known malignancies after multimodal treatments.

### Influence of catheter orientation with respect to WM fibers on drug distribution

This study’s third distinctive feature is the analysis of different catheter positions during in vivo CED procedures, performed parallel or orthogonal to WM tracts. Quantitative evaluations showed that different catheter orientations with respect to the principal axis of the WM bundles influence the initial drug distribution after the CED ceases. At the first imaging TP after stopping CED, infusions performed parallel to CST fibers are significantly more elongated along **λ**1 than those performed orthogonal to the CST, as demonstrated by the computation of EV and Ω_ε1. The interplay between the catheter insertion and the microstructural features of tightly packed axonal bundles is the most likely determining factor of this experimental finding. When the catheter is parallel to the fibers, the infusion direction is aligned to the microstructural constraints that, thus, facilitate the bolus distribution along the principal axis of the WM bundle. When the catheter is perpendicular to the fibers, the infusion direction collides with the brain tissue features, which, in turn, hamper the bolus distribution along the WM course.

Moreover, by observing the shape of the boluses at consecutive imaging TPs, we observed that the infusions parallel to WM are already driven along the main fiber direction at the first TP, continuing to expand preferentially along ε1 over time. This behavior is less evident in infusions orthogonal to WM since in the orthogonal infusion cases, the axons act as obstacles that prevent the infusion penetration along the catheter direction, although those also show a superior elongation trend along ε1. Superimposable results are obtained by comparing the reciprocal evolution of **Ω**_ε1, **Ω**_ε2 and **Ω**_ε3 of infusions parallel or orthogonal to CST fibers, where the difference between **Ω**_ε1 and the other two angles increases over time. Again, this can be explained by the diversion of the infusate in the orthogonal infusion scenario. Specifically, at the very early TPs, flow occurs smoothly in both directions, resulting in a relatively isotropic infusate distribution. After some time, when most of the infused solution has diverted to the direction parallel to the axons, the infusate becomes elongated, leading to a pronounced anisotropic distribution.

The tendency of molecules to preferentially flow lined up with brain axons has been already reported in the literature, both resulting from mathematical model computations (28, 55) and from complex experiments on fresh mammalian brains (56). In particular, the results of Jamal et al. on ex-vivo ovine brain specimens are consistent with our findings since they measured higher hydraulic permeability of brain tissue when axons were aligned to the injection direction and lower hydraulic permeability when axons were orthogonal to the fluid (56). However, our comprehensive demonstration of this phenomenon in living mammals is unique, as it precisely tuned the orientation of catheter insertion in vivo, matching mathematical modeling and experimental findings in a pre-clinical scenario.

This work demonstrated that the final infusate distribution within brain tissue depends on complex intermingled mechanisms: the catheter direction plays a key role especially at the first TPs after stopping CED, then the influence of intrinsic tissue anisotropy prevails. The parallel modelling investigation performed by the team (53) has also demonstrated the importance of the measurement’s framework proposed here to enable a first comprehensive direct validation of advanced multiscale numerical simulations to predict the outcome of CED procedures using in vivo experiments.

### Future perspectives and limitations

This work is predominantly centered on the non-invasive DTI-based evaluation of the brain tissue microstructure to plan and tune CED procedures. However, it lacks the histopathological assessment of infusate distributions. Several studies reported only imaging investigations of drug delivery without any further exploration of tissue features, essentially characterizing CED consequences with sophisticated MRI analyses (39, 41–43, 57–59). Despite in this study we injected a gadolinium-based solution, histopathological evaluations of tissue specimens are particularly beneficial when a potentially toxic drug is delivered within the brain, as they allow to assess the drug’s effects on the microenvironment (60, 61), fiber demyelination (62), changes of vessel permeability and possible astrocyte reactions (38, 40). Another valuable aspect of histology would be assessing the catheter-induced tissue damages (37, 63), especially if innovative delivery-platforms are tested in preclinical settings. In this regard, future experiments, including comprehensive tissue evaluations should be carried on for describing these effects with the newly engineered flexible catheter for minimally invasive neurosurgery (64, 65). We employed animals with no evidence of neurological disorders or pathological imaging features. Further proof should be retrieved in pathological conditions, either neoplastic or degenerative, to assess the reproducibility of the current findings and of the resolution of MR-DTI to depict and assess pathological changes. Disease states can have a significant impact on fluid transport, and, in turn, it is needed to understand how transport can affect disease progression. Glioblastoma, for instance, and its microenvironment, are known to determine a regional change of the fluid flow dynamics.

This study also lacked a specific therapeutic compound. The infusate was a gadolinium-based solution. A longer duration of the observation could also be relevant to assess the infusate distribution over a prolonged time. However, the present results could help guide further CED procedures in the ovine models since new insertion strategies and potential needle steering may also be planned, pondering WM tracts’ orientation. Eventually, an accurate prediction of final infusate distribution may pave the way to more successful and standardized CED clinical trials (66). In fact, even if the development of promising biological therapies remains a challenge (67), technical advances in monitoring the extent of drug delivery are needed in order to guarantee safer and more reproducible therapeutic interventions.

### Conclusion

This study integrated MR tractography reconstructions in the pre-operative planning for CED procedures in the ovine model, in vivo, for the first time. We also provided empirical evidence that the brain tissue constraints, such as the intrinsic anisotropy of axonal bundles, profoundly facilitate the distribution of a gadolinium-based solution along the WM fibers’ main axis while hampering it in the perpendicular direction. Conversely, in an isotropic environment, the infusate distribution does not follow a preferential direction. This comprehensive investigation in living sheep provides essential information for further optimization of the CED procedures that should include DTI-based tissue analyses to predict the actual distribution of the delivered therapy more accurately. It was also shown to be essential to improve the predictive capability and obtain critical information and inputs for the most advanced multiscale simulation tools developed to optimise drug delivery by accurately modelling how therapeutic agents are distributed within the brain, leading to more targeted and effective treatments. This study contributed to the clinical translational of the CED of flow-based therapy to ultimately provide new therapeutic approaches for several brain diseases, with a better prediction model and a delivery platform to reach the target of therapeutic interest and monitor the infusion precisely and non-invasively.

## Supporting information

BIORXIV_2025_648763_SupplementaryMaterial

## Acknowledgements

This work has been carried out in the context of the EDEN2020 (Enhanced Delivery Ecosystem for Neurosurgery in 2020) project, that received funding from the European Union’s EU Research and Innovation programme Horizon 2020 under Grant Agreement No. 688279.

The Authors want to acknowledge the invaluable contributions and support of Marcello Cadioli, our beloved colleague and friend, who prematurely passed away before this paper was submitted. His essential role was instrumental in realizing this project, and his boundless enthusiasm, generosity, and dedication to MRI research will always be remembered.

## Author contributions

Antonella Castellano, Marco Riva, Andrea Falini and Ferdinando Rodriguez y Baena designed the experiments. Antonella Castellano, Valentina Pieri, Marco Riva, Riccardo Secoli, Alice Segato, Marco Vidotto, Marco Trovatelli, Stefano Brizzola, Davide Danilo Zani, Giuliano Ravasio and Stefano Galvan performed the experiments in vivo. Antonella Castellano, Valentina Pieri, Alice Segato and Nicolò Pecco performed the imaging trials and analysis. Riccardo Secoli and Elena De Momi contributed to the data analysis. Tian Yuan and Daniele Dini collaborate to data analysis and refinement for model validation. Antonella Castellano took the lead in writing the article. Marco Riva and Lorenzo Bello help in the refinement of the surgical workflow. Ferdinando Rodriguez y Baena and Andrea Falini supervised the work and contributed to the writing of the article. All authors provided critical feedback and helped shape the research, analysis and manuscript.

## Data availability statement

All data produced in the present study are available upon reasonable request to the authors.

