## Supplementary material for "In vivo MRI measurement of microstructural constraints for direct delivery of therapeutics within the brain": BIORXIV_2025_648763_SupplementaryMaterial

### Supplementary Methods and Tables

**Supplementary Table 1: Criteria to define if the infusate is inside or outside the WM.** This table describes in detail how the two criteria were applied, better clarifying what is summarized in Table1. **A)** Mean and standard deviation of FA values corresponding CSTs location were calculated ($^{CST}FA_{mean}$ = 0.378 and $^{CST}FA_{std}$ = 0.026) as well as mean FA values within each bolus at each timepoint ($^{bolus}FA_{mean}$) (**Supplementary Table1**). Both these steps were carried out by multiplying the FA map times the corresponding binary mask of CSTs and boluses, as explained in algorithm 1 in Supplementary Material, of the same sheep. Next, non-zero voxels of the outputs were averaged by means of 'fslmaths' tool of FSL. The $^{bolus}FA_{mean}$ value was compared to the corresponding $^{CST}FA_{mean}$: if the first differs from the second of more than ± 2* $^{CST}FA_{std}$, the bolus was considered partially outside WM fibers, as in the case of the three infusions marked with (#). **B)** Principal components of the boluses were evaluated with their respective eigenvalues ($\lambda_1$, $\lambda_2$, $\lambda_3$) through PCA (principal component analysis). PCA allowed computing the main eigenvector ($\varepsilon_1$) and its relative eigenvalue ($\lambda_1$) of each bolus. Next, the second and third main directions ($\lambda_2$, $\lambda_3$) were defined as the mutual perpendicular directions. Explained Variance (EV) a is scalar value between 0 and 1, and in this context, it represented the shape of boluses. In particular, a value of 0.33 indicates perfect spheroid shape, while bigger value reflects an ellipsoid shape. EV between $\lambda_1$ and $\lambda_2$, $\lambda_3$ was computed (Equation 1) for each bolus at the first TP: if it resulted inferior to 0.4, the bolus was considered to extend outside WM fibers, as in the case of one infusion (##).

$$EV = \frac{\max(\lambda_1, \lambda_2, \lambda_3)}{\sum_{i=1}^{3} \lambda_i} \quad (1)$$

| | | A) $^{bolus}FA_{mean} = {}^{CST}FA_{mean}$ ± 2SD | | | | | B) EV of $\lambda_1$ > 0.4 |
|---|---|---|---|---|---|---|---|
| **Sheep n°** | **Targeted Hemisphere** | $^{CST}FA_{mean}$ | $^{bolus}FA_{mean}$ | | | | **EV 1st TP** |
| | | | **1st TP** | **2nd TP** | **3rd TP** | **4th TP** | |
| **SHEEP 01** | L # | 0.353010 | 0.2911067 | 0.3039974 | 0.3237782 | -- | -- |
| | R | | 0.3635829 | -- | -- | -- | 0.4838 |
| **SHEEP 02** | L | 0.409366 | 0.5241120 | 0.5194404 | -- | -- | 0.5431 |
| | R | | 0.4691851 | 0.4705228 | 0.4758378 | -- | 0.5563 |
| **SHEEP 03** | L | 0.351904 | 0.4024021 | 0.3804867 | 0.351904 | -- | 0.5368 |
| | R | | 0.4482630 | -- | -- | -- | 0.5508 |
| **SHEEP 04** | L | 0.392805 | 0.5723356 | 0.5293016 | -- | -- | 0.6686 |
| | R # | | 0.3040846 | 0.3150424 | -- | -- | -- |
| **SHEEP 05** | L | 0.395656 | 0.4341159 | 0.4178370 | 0.4148682 | -- | 0.5207 |
| | R | | 0.4163692 | 0.4119800 | 0.4109464 | -- | 0.4675 |
| **SHEEP 06** | L ## | 0.399120 | 0.4244288 | 0.4229526 | 0.4236349 | -- | 0.3801 |
| | R | | 0.3561724 | 0.3621433 | 0.3582588 | -- | 0.4502 |
| **SHEEP 07** | L # | 0.348684 | 0.2505788 | 0.2577671 | 0.2598109 | -- | -- |
| | R | | 0.5254956 | 0.5063998 | 0.5005979 | 0.4995024 | 0.4982 |

**Supplementary Table 2: Numerical data describing the impact of microstructural constraints on diffusion outside WM fibers.** This table reports the data at the basis of graphs in Figure 3. Length of the bolus in the direction of $\varepsilon_1$, $\varepsilon_2$ and $\varepsilon_3$ (L) were computed for each of the four boluses infused outside WM fibers, both at the first and last timepoint (TP) of T1w MRI images acquired after the infusions.

| Infusions OUTSIDE the WM, considering together infusions parallel and orthogonal to CST fibers | | | | | |
|---|---|---|---|---|---|
| **Combination of Parameters** | **Minimum** | **25% Percentile** | **Median** | **75% Percentile** | **Maximum** |
| **$L\_\varepsilon_1$ 1$^{st}$ TP [mm]** | 5.72 | 5.735 | 5.835 | 7 | 7.37 |
| **$L\_\varepsilon_2$ 1$^{st}$ TP [mm]** | 4.2 | 4.263 | 4.74 | 5.488 | 5.64 |
| **$L\_\varepsilon_3$ 1$^{st}$ TP [mm]** | 4.94 | 4.94 | 5.035 | 5.175 | 5.19 |
| **$L\_\varepsilon_1$ last TP [mm]** | 6.12 | 6.373 | 7.2 | 7.84 | 8.03 |
| **$L\_\varepsilon_2$ last TP [mm]** | 4.4 | 4.593 | 5.58 | 6.38 | 6.51 |
| **$L\_\varepsilon_3$ last TP [mm]** | 5.15 | 5.345 | 5.98 | 6.465 | 6.61 |

**Supplementary Table 3: Numerical data describing the impact of catheter orientation on diffusion inside WM fibers.** This table reports the data at the basis of graphs in Figure 5 and 6. Length of the bolus in the direction of $\varepsilon_1$, $\varepsilon_2$ and $\varepsilon_3$ (L) and angle between ε_max_bolus and $\varepsilon_1$ (Ω) were computed for each of the ten boluses infused inside WM fibers, both at the first and last timepoint (TP) of T1w MRI images acquired after the infusions.

| A | Infusions inside the WM, performed with the catheter placed PARALLEL to CST fibers | | | | | |
|---|---|---|---|---|---|---|
| | **Combination of Parameters** | **Minimum** | **25% Percentile** | **Median** | **75% Percentile** | **Maximum** |
| | **$L\_\varepsilon_1$ $1^{st}$ TP [mm]** | 6.29 | 6.545 | 7.55 | 7.8 | 8.01 |
| | **$L\_\varepsilon_2$ $1^{st}$ TP [mm]** | 4.14 | 4.868 | 5.49 | 6.13 | 6.49 |
| | **$L\_\varepsilon_3$ $1^{st}$ TP [mm]** | 4.13 | 4.798 | 5.63 | 5.943 | 6.07 |
| | **$L\_\varepsilon_1$ last TP [mm]** | 6.79 | 7.263 | 7.76 | 9.72 | 9.72 |
| | **$L\_\varepsilon_2$ last TP [mm]** | 4.34 | 5.233 | 6.325 | 7.55 | 7.55 |
| | **$L\_\varepsilon_3$ last TP [mm]** | 4.41 | 4.868 | 6.525 | 6.81 | 6.81 |
| | **$\Omega\_\varepsilon_1$ $1^{st}$ TP [°]** | 7.88 | 16.06 | 25.43 | 46.41 | 56.48 |
| | **$\Omega\_\varepsilon_2$ $1^{st}$ TP [°]** | 60.56 | 65.84 | 78.56 | 83.3 | 87.77 |
| | **$\Omega\_\varepsilon_3$ $1^{st}$ TP [°]** | 74.69 | 75.27 | 80.90 | 87.89 | 89.51 |
| | **$\Omega\_\varepsilon_1$ last TP [°]** | 7.88 | 18.45 | 26.42 | 42.53 | 52.14 |
| | **$\Omega\_\varepsilon_2$ last TP [°]** | 55.90 | 57.13 | 71.44 | 81.86 | 84.63 |
| | **$\Omega\_\varepsilon_3$ last TP [°]** | 69.35 | 70.53 | 79.09 | 87.89 | 89.52 |
| **B** | **Infusions inside the WM, performed with the catheter placed ORTHOGONAL to CST fibers** | | | | | |
| | **Combination of Parameters** | **Minimum** | **25% Percentile** | **Median** | **75% Percentile** | **Maximum** |
| | **$L\_\varepsilon_1$ $1^{st}$ TP [mm]** | 5.7 | 5.815 | 6.22 | 6.678 | 6.81 |
| | **$L\_\varepsilon_2$ $1^{st}$ TP [mm]** | 4.72 | 5.025 | 6.075 | 6.405 | 6.47 |
| | **$L\_\varepsilon_3$ $1^{st}$ TP [mm]** | 4.39 | 4.45 | 5.215 | 5.988 | 6.05 |
| | **$L\_\varepsilon_1$ last TP [mm]** | 6.72 | 6.81 | 7.355 | 7.99 | 8.12 |
| | **$L\_\varepsilon_2$ last TP [mm]** | 5.97 | 6.125 | 6.77 | 6.988 | 7 |
| | **$L\_\varepsilon_3$ last TP [mm]** | 5.07 | 5.16 | 5.935 | 6.62 | 6.68 |
| | **$\Omega\_\varepsilon_1$ $1^{st}$ TP [°]** | 16.61 | 23.43 | 54.58 | 72.96 | 75.52 |
| | **$\Omega\_\varepsilon_2$ $1^{st}$ TP [°]** | 32.36 | 36.51 | 56.17 | 77.75 | 82.82 |
| | **$\Omega\_\varepsilon_3$ $1^{st}$ TP [°]** | 41.50 | 44.61 | 61.8 | 81.32 | 85.2 |
| | **$\Omega\_\varepsilon_1$ last TP [°]** | 14.41 | 15.75 | 44.59 | 62.77 | 68.6 |
| | **$\Omega\_\varepsilon_2$ last TP [°]** | 35.5 | 44.85 | 62.01 | 85.17 | 86.34 |
| | **$\Omega\_\varepsilon_3$ last TP [°]** | 44.64 | 50.34 | 69.54 | 87.41 | 87.74 |

**Supplementary Table 4: Region-Growing algorithm for infusion segmentation.**

An automatic region-growing (RG) algorithm was created in Matlab (vR2019a, Mathworks) to extract the binary masks of the infused boluses for each TP. This process led to optimal segmentations that corresponded to the observed boluses edge. In this study, the maximum voxel intensity of each bolus was found through ITK-SNAP (http://www.itksnap.org) by creating a 3D-square-mask of size 10 voxels that could widely incorporate the whole bolus in the first post-infusion volume for each sheep (**Figure 1F**). The maximum voxel intensity value within the mask was then extracted by using the "fslmaths" tool of FSL (**Table 4**). The relative minimum voxel intensity value of the bolus was detected by visual inspection by a board-certified neuroradiologist. These two values allowed computing the signal intensity decay as a percentage between bolus maximum voxel intensity and its relative minimum (**Table 4**). Mean and standard deviation of percentage values were evaluated (33% ± 3.32) and applied to all maxima, obtaining the growth thresholds for the automatic segmentation algorithm. The maximum voxel intensity value within the mask was taken as the upper value for the seed threshold. The algorithm was instructed to limit the segmentations only to those voxels that displayed an intensity higher than 33% of the maximum gadolinium intensity. Thus, until a 66% of intensity-decay, voxels were still automatically considered as part of the infused bolus mask, while voxels less intense than the growth-threshold were not considered.

| Sheep n° | Targeted Hemisphere | ROI max voxel intensity | ROI min voxel intensity | Signal intensity Decay (%) |
|---|---|---|---|---|
| **SHEEP 01** | L | 5.69 | 1.80 | 31.63 |
| | R | 7.89 | 2.83 | 35.87 |
| **SHEEP 02** | L | 6.36 | 2.54 | 39.94 |
| | R | 5.17 | 1.48 | 28.63 |
| **SHEEP 03** | L | 5.44 | 1.69 | 31.07 |
| | R | 5.17 | 1.57 | 30.37 |
| **SHEEP 04** | L | 5.14 | 1.47 | 28.60 |
| | R | 8.00 | 3.05 | 38.13 |
| **SHEEP 05** | L | 5.61 | 1.85 | 32.98 |
| | R | 5.52 | 1.90 | 34.42 |
| **SHEEP 06** | L | 7.31 | 2.54 | 34.75 |
| | R | 5.55 | 1.72 | 30.99 |
| **SHEEP 07** | L | 5.85 | 1.91 | 32.65 |
| | R | 5.91 | 1.89 | 31.98 |

**Algorithm 1** Computation of the diffusion signal with the tensor model

```
Input: 3DTI_FFE volume dti, initialisation vectors bvals and bvec, bolus geometry bolusCord
tensorFit ← computeImageTensor(dti, bvals, bvecs)
FA_dti ← tensorFit[0]
ε_dti ← tensorFit[1]
λ_dti ← tensorFit[2]
ε_tot ← 0
FA_tot ← 0
for voxel in bolusCord do
    components ← PCA(voxel, FA_dti, ε_dti, λ_dti)
    ε_1 ← components[0]
    ε_2 ← components[1]
    ε_3 ← components[2]
    λ_1 ← components[3]
    λ_2 ← components[4]
    λ_3 ← components[5]
    FA ← components[6]
    ε_tot ← ε_tot + ε_1
    FA_tot ← FA_tot + FA
end for
bolus_mean ← mean(ε_tot, FA_tot)
ε_max_bolus ← bolus_mean[0]
FA_mean ← bolus_mean[1]
return ε_max_bolus, FA_mean
```

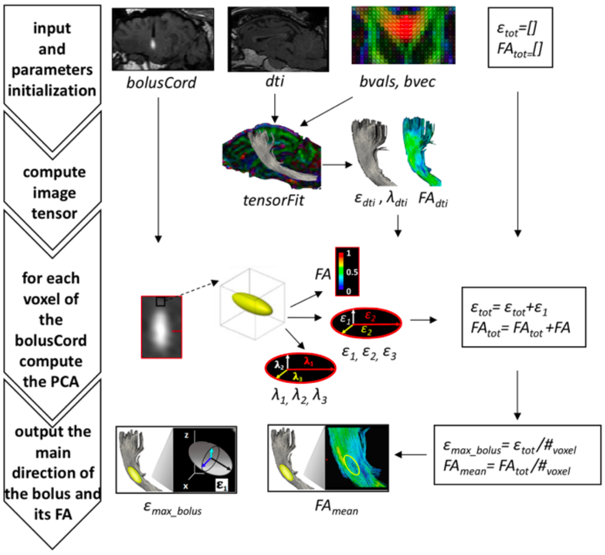


**Algorithm 1** describes the steps to compute the Principal component analysis PCA relative to the maximal direction of the bolus ($\varepsilon\_max\_bolus$), the average FA of the infusate bolus ($FA_{mean}$) as well as to extract the DTI-derived eigenvectors ($\varepsilon_1$, $\varepsilon_2$, $\varepsilon_3$) and eigenvalues ($\lambda_1$, $\lambda_2$, $\lambda_3$). These parameters were calculated for all the fourteen infusions taking in input the Post-infusion 3DTI_FFE scan, the initialization parameters of *bvals* and *bvec* and the coordinates of the geometry of the bolus, *bolusCord*. The algorithm uses the information about the gradients *bvals* and *bvec* to fit the model on the input *dti* images. The *computeImageTensor* method creates a *tensorFit* object which contains the fitting parameters and other attributes of the model. In this way, we can calculate the fractional anisotropy (FA) from the eigenvalues of the tensor. $FA_{dti}$ is used to characterize the degree to which the distribution of diffusion in a voxel is directional. The algorithm subsequently finds the eigenvectors $\varepsilon_{dti}$ and the eigenvalues $\lambda_{dti}$ of the input image *dti*.

The algorithm then loops on the coordinates of the geometry of the bolus *bolusCord* to compute the PCA for each voxel to obtain the primary ($\varepsilon_1$), secondary ($\varepsilon_2$) and minimal ($\varepsilon_3$) component of the *dti* voxel calculated taking into consideration the corresponding maximum ($\lambda_1$) medium ($\lambda_2$) and minimum ($\lambda_3$) eigenvalue weighted for the *FA* of the corresponding voxel.

Finally, the algorithm returns the maximal direction of the bolus, $\varepsilon\_max\_bolus$, calculated as the average of the primary components $\varepsilon_1$, of each voxel and $FA_{mean}$ calculate as the average of the *FA* of each voxel.
